# Small-angle neutron scattering solution structures of NADPH-dependent sulfite reductase

**DOI:** 10.1101/2020.12.08.415968

**Authors:** Daniel T. Murray, Kevin L. Weiss, Christopher B. Stanley, Gergely Nagy, M. Elizabeth Stroupe

**Affiliations:** Department of Biological Science and Institute of Molecular Biophysics, Florida State University, Tallahassee, Florida, 32306, USA; Neutron Scattering Division, Oak Ridge National Laboratory, Oak Ridge, Tennessee, 37830, USA

**Keywords:** analytical ultracentrifugation, assimilatory sulfite reductase, electron transfer, oxidoreductase, small-angle neutron scattering

## Abstract

Sulfite reductase (SiR), a dodecameric complex of flavoprotein reductase subunits (SiRFP) and hemoprotein oxidase subunits (SiRHP), reduces sulfur reduction for biomass incorporation. Electron transfer within SiR requires intra- and inter-subunit interactions that are mediated by the relative position of each protein, governed by flexible domain movements. Using small-angle neutron scattering, we report the first solution structures of SiR heterodimers containing a single copy of each subunit. These structures show how the subunits bind and how both subunit binding and oxidation state impact SiRFP’s conformation. Neutron contrast matching experiments on selectively deuterated heterodimers allow us to define the contribution of each subunit to the solution scattering. SiRHP binding induces a change in the position of SiRFP’s flavodoxin-like domain relative to its ferredoxin-NADP^+^ reductase domain while compacting SiRHP’s N-terminus. Reduction of SiRFP leads to a more open structure relative to its oxidized state, re-positioning SiRFP’s N-terminal flavodoxin-like domain towards the SiRHP binding position. These structures show, for the first time, how both SiRHP binding to, and reduction of, SiRFP positions SiRFP for electron transfer between the subunits.

## INTRODUCTION

Multi-subunit oxidoreductase enzymes catalyze electron transfer reduction-oxidation (redox) reactions that drive energy flux in cells. A conserved class of NADPH-dependent diflavin reductases funnel electrons from NADPH through flavin adenine dinucleotide (FAD) and flavin mononucleotide (FMN) cofactors to diverse oxidases that are often metalloenzymes. Assimilatory NADPH-dependent sulfite reductase (SiR) is a member of this class of enzymes and is responsible for the six-electron reduction of sulfite (SO_3_^2-^) to sulfide (S^2-^) for incorporation into sulfur-containing biomolecules (Siegel *et al*, 1973). These SiRs are dodecameric, with a uniquely octameric diflavin reductase (SiRFP, the α subunit) and four copies of a siroheme- and Fe_4_S_4_-containing hemoprotein (SiRHP, the β subunit) (Murphy *et al*, 1973; Siegel & Davis, 1974; Siegel *et al*, 1974).

SiR’s α_8_β_4_ dodecameric assembly sets it apart from other members of this class of diflavin reductase-dependent enzymes like cytochrome P450 reductase (CPR), nitric oxide synthase (NOS), cytochrome P102, and methionine synthase (Campbell *et al*, 2014; Olteanu & Banerjee, 2001; Xia *et al*, 2011b; Zhang *et al*, 2018). Based on what is known about the well-studied CPR (Freeman *et al*, 2018; Freeman *et al*, 2017), electron transfer within the SiR diflavin reductase subunit is hypothesized to work through a redox-sensitive conformational change between two domains that are separated by a flexible hinge. Specifically, the FMN-binding domain, which is homologous to small flavodoxins (Fld), moves close to the NADPH-reduced FAD, which binds to a ferredoxin-NADP^+^ reductase (FNR) domain, to itself become reduced (Huang *et al*, 2013; Iyanagi *et al*, 2012; Wang *et al*, 1997). Upon NADP^+^ release, the Fld domain swivels back out to pass the electrons to its oxidase partner (Hamdane *et al*, 2009; Xia *et al*, 2011a). In CPR, the opening of the Fld domain creates the binding site for the heme-containing oxidase (Im & Waskell, 2011). In NOS, where the two subunits are expressed in a single polypeptide that dimerizes, this conformational change allows the reductase from one subunit to interact with the heme-binding oxidase domain from the partner (Campbell *et al*., 2014; Haque *et al*, 2018). The absence of any identified structure of the bound SiR subunits leaves a gap in our understanding of how intersubunit interactions and domain motions govern electron transfer in SiR.

Another aspect of SiR that sets it apart from analogous systems is the way in which SiRFP and SiRHP interact, through interactions between SiRHP’s N-terminus and a position on SiRFP’s FNR domain that is far from the Fld domain (Askenasy *et al*, 2018; Askenasy *et al*, 2015). This interaction is counterintuitive because the Fld domain must interact with SiRHP to pass electrons to its siroheme-Fe_4_S_4_ cofactors that funnel the electrons to the siroheme-bound substrate. Nevertheless, a separate, transient interaction between SiRFP’s Fld domain and a tightly-bound SiRHP is sufficient for SiR activity because a heterodimeric complex between a monomeric form of SiRFP and full-length SiRHP is active for SO_3_^2-^ reduction, albeit at reduced efficiency (Figures 1A-C, Gruez *et al*, 2000; Zeghouf *et al*., 2000). SiR’s capability for electron transfer is further affected if the flexible linker mediating domain movements of SiRFP is truncated, reducing activity in the heterodimer but not the dodecamer (Tavolieri *et al*, 2019). These results suggest a model in which the dodecamer transfers electrons through multiple pathways, first from an NADPH to the FAD and on to the FMN cofactor, either within a single SiRFP (intramolecular transfer) or from the FAD of one subunit to the FMN of an adjacent molecule (intermolecular transfer), and then either to a tightly bound SiRHP (in *cis*, Figure 1D) or to a SiRHP on an adjacent SiRFP subunit (in *trans*, Figure 1D). This redundancy in electron donor/acceptor pairing could explain SiR’s ability to catalyze high-volume electron transfer without releasing partially-reduced intermediates (Hsieh *et al*, 2010; Lancaster, 2018; Mirts *et al*, 2018; Oliveira *et al*, 2011).

**Figure 1:**
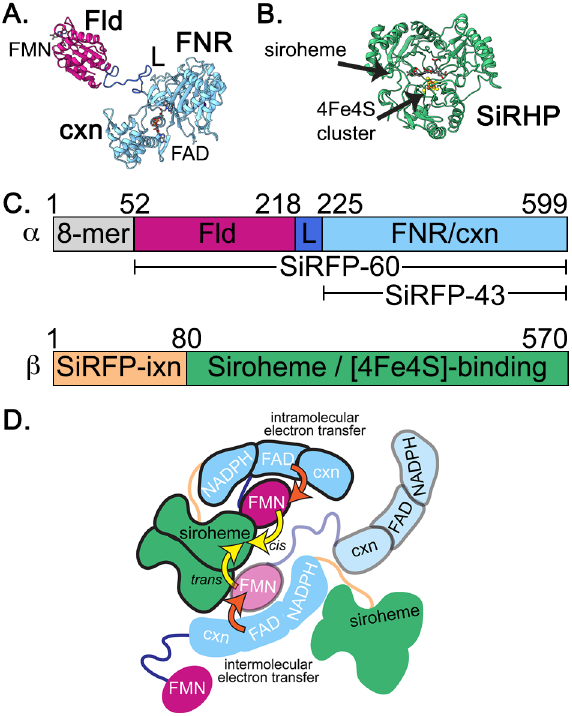
SiR domain composition and structure. **A**. SiRFP, the α subunit (PDB 6EFV (Tavolieri *et al*., 2019); 8-mer is octamerization domain; L is flexible linker; Fld is flavodoxin-like domain; FNR/cxn are ferredoxin-NADP+ reductase and connection domains). **B**. SiRHP, the β subunit (PDB 1AOP (Crane *et al*., 1995); SiRFP-ixn is the N-terminal SiRFP-interaction domain). This color scheme for SiRFP domains and SiRHP are used throughout the manuscript. **C**. Domain composition of SiRFP and SiRHP, colored as in **A**. and **B. D**. Redundancy in possible electron transfer pathways include intra-versus inter-molecular transfer within SiRFP (orange arrows) and *cis* versus *trans* electron transfer between SiRFP and SiRHP (yellow arrows). Bold letters show the domain nomenclature while non-bold letters label the cofactors. Tightly-bound SiRFP/SiRHP dimers are outlined in the same way, either black, gray, or none.

In this study, we used small-angle neutron scattering (SANS), selective deuteration, solvent contrast variation, anaerobic reductions, and analytical ultracentrifugation (AUC) to probe the effects of altering the redox state and subunit binding on heterodimers of a monomeric SiRFP and its SiRHP partner. AUC on SiR heterodimers revealed sedimentation coefficients consistent with their respective oligomeric states as *in vitro* reconstituted specimen. SANS was used to measure hydrodynamic parameters and uncover the first-ever solution structures of these heterodimers. Through the use of selective deuteration and neutron contrast variation, the isolated scattering components of SiR heterodimers illuminated the contributions from each subunit to their complexes, informing our understanding of this multisubunit enzyme. Anaerobically reduced SiRFP variants were similarly measured by SANS and showed movements between the Fld and FNR domains. Together, these observations help explain the nature of its complex assembly as well as its capacity for electron transfer.

## RESULTS

### *In vitro* reconstitution of SiR complexes and their analysis by AUC

SiR subunits bind one another through interactions between SiRFP’s FNR domain and SiRHP’s N-terminus (Askenasy *et al*., 2018; Askenasy *et al*., 2015) but the structure of the resulting complex is unknown. The monomeric SiRFP variants used in this study include one in which the 52 N-terminal amino acids are removed to prevent it from octamerization, retaining the Fld and FNR/connection domains but resulting in a 60 kDa monomer^*42*^(SiRFP-60, Figure 1C). Additional internal truncation of the linker between the Fld and FNR domains is required to immobilize the monomer for structure determination (Tavolieri *et al*., 2019), SiRFP-60-Δ. An additional variant consists solely of its C-terminal, 43 kDa FNR/connection domains (Covès *et al*, 1999; Zeghouf *et al*., 2000, SiRFP-43), representing a minimal, inactive version of SiRFP (Figure 1C). Full-length SiRHP was used in all studies because when its N-terminus is removed it can no longer bind SiRFP (Askenasy *et al*., 2015); however, those amino acids are not present in the X-ray crystal structure (Crane *et al*, 1995, Figure 1B).

SiRFP/SiRHP complexes were formed by 1:1 *in vitro* reconstitution. Each subunit and their resulting heterodimers were subjected to sedimentation velocity AUC to confirm that the subunits assembled free of residual monomer (Figure 2). Each sample exhibited sedimentation coefficient (S) envelopes with single peaks proportional to their molecular weight. The lowest molecular weight fragment of the series, SiRFP-43, sedimented with 3.2 S. SiRFP-60 sedimented at 3.9 S and the 64 kDa SiRHP sedimented at 4.7 S. Additionally, SiRFP-43/SiRHP and SiRFP-60/SiRHP sedimented at 5.5 S and 6.1 S, respectively, and both heterodimers showed distributions with clearly defined, single peaks. These envelopes are consistent with monodisperse specimen suitable for solution scattering experiments, a prerequisite for accurately interpreting scattering data.

**Figure 2:**
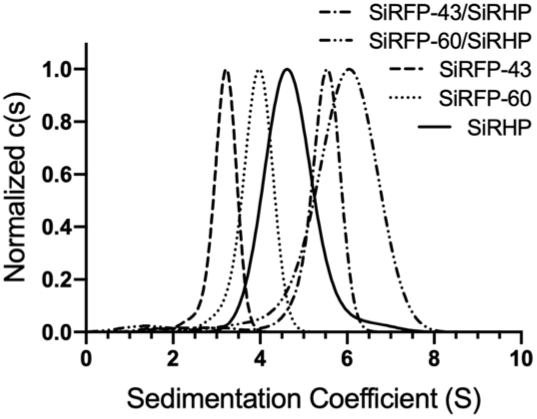
SiR heterodimers are monodispersed. Sedimentation velocity AUC on SiR monomers (SiRFP-43, SiRFP-60, and SiRHP) and heterodimers (SiRFP-43/SiRHP and SiRFP-60/SiRHP) show sedimentation coefficient envelopes are dominated by single peaks, indicative of monodisperse specimen.

### Solution structures of SiR heterodimers

Although structures of the monomeric SiR subunits are known from X-ray crystallography (Crane *et al*., 1995; Gruez *et al*., 2000; Tavolieri *et al*., 2019), no structures exist for higher-order complexes. Consequently, we do not know how the subunits interact such that there is a tight-binding, structural interface independent of the transient, functional interface that mediates electron transfer (Askenasy *et al*., 2018; Askenasy *et al*., 2015). Additionally, we do not know if subunit assembly affects the relative domain orientations of the conformationally dynamic SiRFP. We therefore measured neutron scattering of 1:1 SiR heterodimers.

Scattering of the minimal SiRFP-43/SiRHP heterodimer particle resulted in a radius of gyration, *R*_*g*_, of 32.3 Å and a maximum linear distance, *D*_*max*_, of 99 Å (Figure 3A and Table 1). The bimodal distance distribution function, *P(r)*, of SiRFP-43/SiRHP’s scattering profile is consistent with a multi-domain complex, as expected (Figure 3B). *Ab initio* modeling produced an envelope function that reflects this feature with a real space correlation coefficient (RSC) of 0.86 to an estimated 38 Å resolution with good convergence of the *χ*^2^, *R*_*g*_, and support volume (Figures 3C,S1A, and Table 2). The theoretical scattering curve calculated from the envelope matches the experimental data with low residuals (Figure S1A). The curved shape of one domain was consistent with the dimensions of SiRFP-43, whereas the globular shape of the other domain was consistent with the dimensions of SiRHP, so we placed atomic models accordingly (Figure 3C). Maps calculated from the atomic models and filtered to 20 Å-resolution fit with a correlation coefficient (CC) of 0.71, calculated in ChimeraX (Goddard *et al*, 2018) Table 2). Further, the theoretical scattering curve calculated from the docked X-ray crystal structures of the individual subunits is consistent with the experimental scattering (Figure S2A). To test our assignment of the curved FNR domain, we also measured the scattering of SiRFP-43, which yielded an *R*_*g*_ of 24.7 Å and *D*_*max*_ of 83 Å (Table 1), and whose envelope revealed the expected curve shape, consistent with SiRFP’s FNR/connection domains (Figure S3A).

**Table 1:** Hydrodynamic parameters of SiR subunits and reconstituted dimers determined by SANS.

| Table 1: Hydrodynamic parameters of sulfite reductase |  |  |  |  |  |  |
| --- | --- | --- | --- | --- | --- | --- |
| Sample | D<sub>2</sub>O (%) | Domain/protein composition | Mass (kDa) | Oligomeric state | $R_g$ (Å) | $D_{max}$ (Å) |
| SIRFP-43/SIRHP | 90 | FNR/SIRHP | 107 | Dimer ($\alpha\beta$) | 32.3 | 99 |
| SIRFP-43 | 90 | FNR | 43 | Monomer ($\alpha$) | 24.7 | 83 |
| SIRFP-60/SIRHP | 90 | Fld-linker-FNR/SIRHP | 124 | Dimer ($\alpha\beta$) | 38.0 | 134 |
| SIRFP-60-Δ/SIRHP | 90 | Fld-ΔAAPSQS-FNR/SIRHP | 123 | Dimer ($\alpha\beta$) | 37.8 | 132 |
| SIRFP-60/D-SIRHP* | 86 | Fld-linker-FNR/D-SIRHP | 124 | Dimer ($\alpha\beta$) | 33.1 | 124 |
| SIRFP-60 | 90 | Fld-linker-FNR | 60 | Monomer ($\alpha$) | 32.2 | 113 |
| SIRFP-43/D-SIRHP* | 86 | FNR/D-SIRHP | 107 | Dimer ($\alpha\beta$) | 25.2 | 73 |
| SIRFP-60-Δ/D-SIRHP* | 86 | Fld-ΔAAPSQS-FNR/D-SIRHP | 123 | Dimer ($\alpha\beta$) | 33.5 | 123 |
| SIRFP-60*/D-SIRHP | 41 | Fld-linker-FNR/D-SIRHP | 124 | Dimer ($\alpha\beta$) | 24.1 | 66 |
| D-SIRHP | 41 | D-SIRHP | 64 | Monomer ($\beta$) | 25.9 | 75 |
| SIRFP-43*/D-SIRHP | 41 | FNR/D-SIRHP | 107 | Dimer ($\alpha\beta$) | 23.7 | 65 |
| SIRFP-60-Δ*/D-SIRHP | 41 | Fld-ΔAAPSQS-FNR/D-SIRHP | 123 | Dimer ($\alpha\beta$) | 23.6 | 68 |
| SIRFP-60 Reduced | 90 | Fld-linker-FNR | 60 | Monomer ($\alpha$) | 33.8 | 122 |
| SIRFP-60-Δ Reduced | 90 | Fld-ΔAAPSQS-FNR | 59 | Monomer ($\alpha$) | 33.2 | 114 |

\*Matched out component

**Table 2:** Real space correlations, Fourier shell correlation resolutions, and correlation coefficients of *ab initio* solution structure models.

| Table 2: Model Validation Statistics |  |  |  |  |
| --- | --- | --- | --- | --- |
| Sample | D<sub>2</sub>O (%) | DENSS |  | CHIMERAX |
|  |  | Real space correlation | Fourier Shell Correlation Resolution (Å) | CC** |
| SIRFP-43/SIRHP | 90 | 0.86 | 38 | 0.71 |
| SIRFP-60/SIRHP | 90 | 0.89 | 38 | 0.76 |
| SIRFP-60/D-SIRHP* | 86 | 0.88 | 31 | 0.71 |
| SIRFP-60 | 90 | 0.89 | 29 | 0.71 |
| SIRFP-60*/D-SIRHP | 41 | 0.95 | 20 | 0.89 |
| D-SIRHP | 41 | 0.86 | 33 | 0.66 |
| SIRFP-60 Reduced | 90 | 0.88 | 28 | 0.80 |

\*Matched out component
\*\* CC describes the fit between docked experimental model from the known coordinates, filtered at 20 Å, and the DENSS envelope function, calculated in ChimeraX

**Figure 3:**
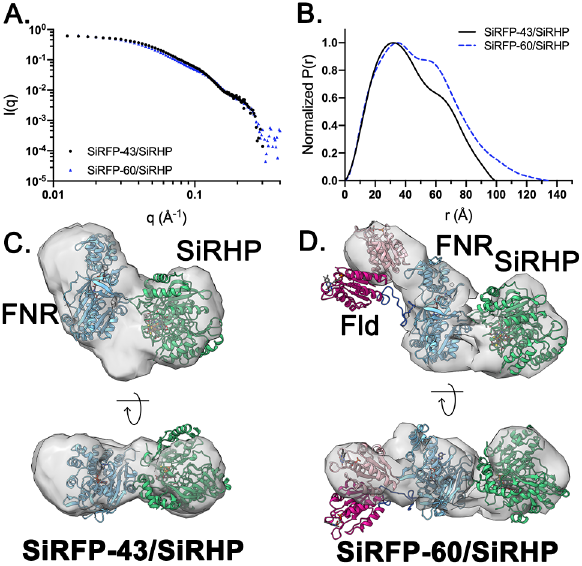
Solution structures of heterodimeric SiR: SiRFP-43/SiRHP and SiRFP-60/SiRHP. **A**. Scattering profiles of SiRFP-43/SiRHP and SiRFP-60/SiRHP show gradual decay into higher *q* regions. **B**. Distance distribution functions for heterodimers’ scattering in **A**. show multicomponent systems. The *ab initio* models (grey density) of SiRFP-43/SiRHP (**C**.) and SiRFP-60/SiRHP (**D**.) with high-resolution structures superimposed (**C**.) SiRFP-43, PDB 1DDG (Gruez *et al*., 2000) and (**D**.) SiRFP-60-Δ, PDB 6EFV (Tavolieri *et al*., 2019)). High-resolution structures are colored as in **Figure 1**. In **D**., the rearranged Fld domain is colored light pink.

The larger SiRFP-60/SiRHP heterodimer scatters with an *R*_*g*_ of 38.0 Å and a *D*_*max*_ of 134 Å (Figure 3A and Table 1). Compared to the *P(r)* calculated from the scattering curve for the SiRFP-43/SiRHP dimer, the *P(r)* calculated from the scattering curve for SiRFP-60/SiRHP shows a pronounced bimodal distribution that gradually tails off, suggesting an elongated structure with greater delineation of the domains than in the SiRFP-43/SiRHP dimer (Figure 3B). *Ab initio* modeling produced a multi-lobed structure with a central curved domain adjacent to a globular domain of similar shape and size to the SiRFP-43/SiRHP dimer (Figures 3C and D). An additional domain with the same dimensions as SiRFP’s N-terminal Fld domain appeared, opposed to the globular domain. The modeled envelope function matches the experimental data well (RSC = 0.89, resolution = 38 Å), with good convergence statistics (Figure S1B and Table 2). The theoretical scattering curve calculated from the envelope matches the experimental data with low residuals (Figure S1B). Further, docking the X-ray crystal structure of SiRFP-60-Δ into the envelope shows that SiRFP-60’s FNR/connection domains fit well into the central curved domain but with a mismatch in the position of the Fld domain that was resolved by a rigid body motion, whereas SiRHP fit into the globular domain as in the SiRFP-43/SiRHP dimer (CC = 0.76, Figure 3D and Table 2). The theoretical scattering curve calculated from the docked X-ray crystal structures of SiRFP-60/SiRHP is consistent with its experimental scattering curve (Figure S2B). We also collected neutron scattering on the dimer formed by assembling SiRFP-60-Δ with SiRHP (SiRFP-60-Δ/SiRHP) as a control to determine if the full-length linker was too flexible to accurately model. The *R*_*g*_, *D*_*max*_, and envelope were nearly identical to those determined for SiRFP-60/SiRHP (Table 1 and Figure S3B).

### Isolated SiRFP-60 scattering within the heterodimer

To confirm our domain assignment in the SiRFP-60/SiRHP heterodimer, we isolated the scattering contribution of each subunit by selectively deuterating SiRHP (D-SiRHP) and manipulating the solvent contrast. The subunit that was matched-out by manipulating the H_2_ O:D_2_ O ratio will be demarked by an asterisk throughout. SANS measurements on SiRFP-60/D-SiRHP* in buffer at D-SiRHP’s contrast match point (CMP, Figure S4) isolated the scattering from SiRFP-60 in the context of its heterodimer structure (Figure 4A). The *R*_*g*_ and *D*_*max*_ of SiRFP-60/D-SiRHP* is larger than the reported values for monomeric SiRFP-60, previously measured by SANS (Tavolieri *et al*., 2019). Specifically, the *R*_*g*_ increased from 32.2 to 33.1 Å and its *D*_*max*_ increased from 113 to 124 Å (Table 1). This manifests in a *P(r)* plot for SiRFP-60/D-SiRHP* whose distribution function decays like an extended molecule, which is reflected in its *ab initio* model (RSC = 0.88, resolution = 31 Å) with good convergence (Figures 4B,C, S1, and Table 2). The extended, multi-lobed shape agrees with our assignment of SiRFP-60’s position in the heterodimer, including the Fld domain repositioned from its place in the X-ray structure of SiRFP-60-Δ and solution structure of SiRFP-60 (Tavolieri *et al*., 2019, CC = 0.71 for both models, Figures 4C,D, and Table 2). Theoretical scattering curves calculated from the envelope function as well as the docked atomic models correlate with the experimental data with low residuals (Figures S1C and S2C).

**Figure 4:**
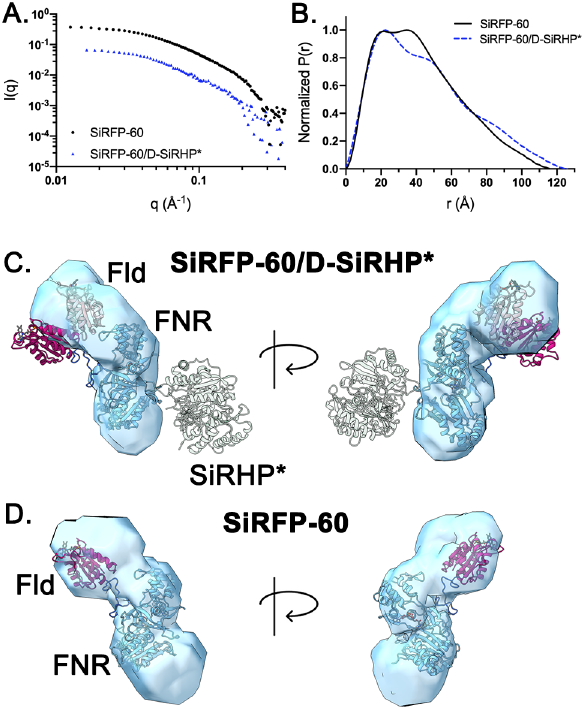
SiRFP-60 exhibits an extended conformation upon binding SiRHP. **A**. Scattering profile of SiRFP-60/D-SiRHP* (D-SiRHP matched-out) and SiRFP-60. **B**. The distance distribution plot of SiRFP-60/D-SiRHP* exhibits a shift in its bilobed peaks relative to the previously reported, oxidized form (Tavolieri *et al*., 2019). **C**. Model (blue density) of SiRFP-60/D-SiRHP* with SiRFP-60-Δ’s crystal structure superimposed, colored as in **Figure 1**. Matched-out D-SiRHP is shown transparently and the rearranged Fld domain is colored light pink. **D**. SiRFP-60’s solution structure, as previously reported (Tavolieri *et al*., 2019).

As a control, we also measured the scattering of SiRFP-43/D-SiRHP* at D-SiRHP’s CMP, which revealed only subtle changes to the *R*_*g*_*/D*_*max*_ and the envelope of SiRHP-bound SiRFP-43 compared to its monomeric form, as expected due to the absence of the Fld domain in SiRFP-43 (Figure S3C and Table 1). In addition, scattering of SiRFP-60-Δ/D-SiRHP* agreed with SiRFP-60/D-SiRHP*, further supporting our interpretation of the Fld domain motion upon SiRHP binding (Figure S3D and Table 1).

### SiRHP undergoes compaction upon binding SiRFP-60

We next measured the contribution of D-SiRHP to the scattering of the SiRFP-60/D-SiRHP heterodimer by preparing SiRFP-60/D-SiRHP in buffers at SiRFP-60’s CMP (Figure 5A). Compared to the scattering of monomeric D-SiRHP, SiRFP-60*/D-SiRHP exhibited a decrease in its *R*_*g*_ from 25.9 to 24.1 Å. *D*_*max*_ values obtained from *P(r)* analyses of D-SiRHP and SiRFP-60*/D-SiRHP reflected this compaction, from 75 to 66 Å, respectively (Figure 5B and Table 1). Solution structures of SiRFP-60*/D-SiRHP and D-SiRHP obtained from *ab initio* modeling also manifest these differences with a rearrangement of density in the models between the monomer and heterodimer (RSC = 0.95 and 0.86 to estimated resolutions of 19.8 and 32.7 Å, respectively, Figures 5C,D, and Table 2). Theoretical scattering curves calculated from the *ab initio* models of SiRFP-60*/D-SiRHP and D-SiRHP both correspond to their respective experimental scattering curves with low residuals, as does the theoretical scattering curve calculated from the atomic model of SiRHP (Crane *et al*., 1995, Figures S1D,E, S2D, and E). The atomic model of SiRHP fits into the volumes representing SiRFP-60*/D-SiRHP or D-SiRHP with a CC of 0.71 or 0.89, respectively (Table 2). As controls, we measured D-SiRHP’s contribution to the scattering of the other heterodimer variants (SiRFP-60-Δ*/D-SiRHP and SiRFP-43*/D-SiRHP), which also showed contraction of D-SiRHP in both the hydrodynamic parameters and modeled envelopes (Figures S3E and F, Table 1).

**Figure 5:**
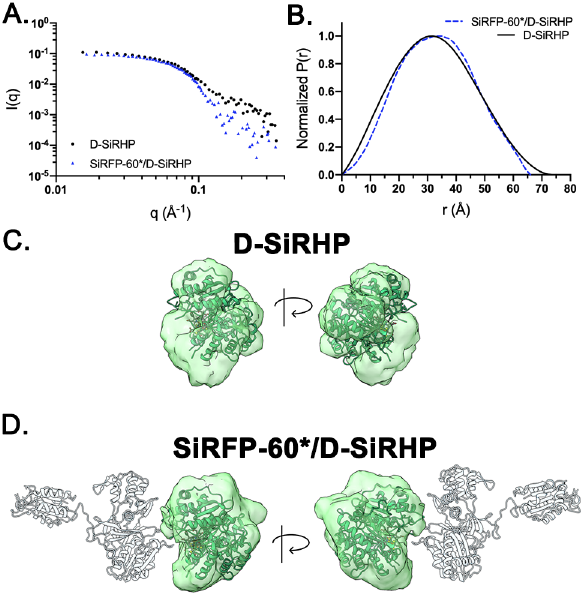
SiRHP undergoes compaction upon binding SiRFP. **A**. Scattering profiles of D-SiRHP and SiRFP-60*/D-SiRHP. **B**. Distance distribution plots for the scattering profiles in **A**. show D-SiRHP adopts a reduced *D*_*max*_ upon binding SiRFP-60. **C**. Models (green density) of D-SiRHP and **D**. SiRFP-60*/D-SiRHP. SiRHP’s crystal structure is superimposed on both models and SiRFP-60 is shown transparently to reflect its matching-out in **D**.

### SiRFP-60’s Fld domain further repositions upon dithionite reduction

By analogy with homologous enzymes like CPR and NOS (Campbell *et al*., 2014; Huang *et al*., 2013), we predicted that SiRFP-60 exhibits redox-dependent domain movements that allow reducing equivalents to pass from the NADPH-reduced FAD to the FMN cofactor. In CPR, the domains open relative to one another upon FMN reduction to allow transfer of reducing equivalents to the heme-containing oxidase. Unlike CPR, however, SiRFP-60 adopts an extended state in solution when it is oxidized (Figures 4A,B, and D (Huang *et al*., 2013; Tavolieri *et al*., 2019)). Therefore, to test the effect of redox state on SiRFP’s structure, SiRFP-60 was reduced with 10 equivalents of sodium dithionite in an anaerobic glovebox before being transferred to a cuvette sealed from atmosphere and measured with SANS.

The scattering profile of reduced SiRFP-60 indicates an open conformation (Figure 6A), even more exaggerated than its oxidized conformation (Tavolieri *et al*., 2019). Compared to the previously reported solution structure of oxidized SiRFP-60 (Figure 4D), 10 Eq-reduced SiRFP-60’s *R*_*g*_ increases from 32.2 to 33.8 Å and the *D*_*max*_ shifts from 113 to 122 Å (Figure 6B and Table 1) whereas the envelope of reduced SiRFP-60 (RSC = 0.88, resolution = 28.1 Å, Table 2) shows its characteristic bent FNR domain and the globular Fld domain in a novel conformation (Figure 6C). The theoretical scattering curve calculated from the *ab initio* model corresponds to its experimental scattering curve with low residuals and the model resulting from the repositioned Fld domain fits the experimental scattering well (CC = 0.80, Figures S1F,S2F, and Table 2). We also measured the scattering of dithionite-reduced SiRFP-60-Δ, which similarly showed extension compared to its oxidized state, albeit not as dramatic as in SiRFP-60 (Figure S3G and Table 1) (Tavolieri *et al*., 2019). Note, this experiment cannot be performed on the heterodimers because oxidized dithionite is a substrate for SiRHP, so the system would be in multiple redox states as it consumes the dithionite (Siegel *et al*., 1974).

**Figure 6:**
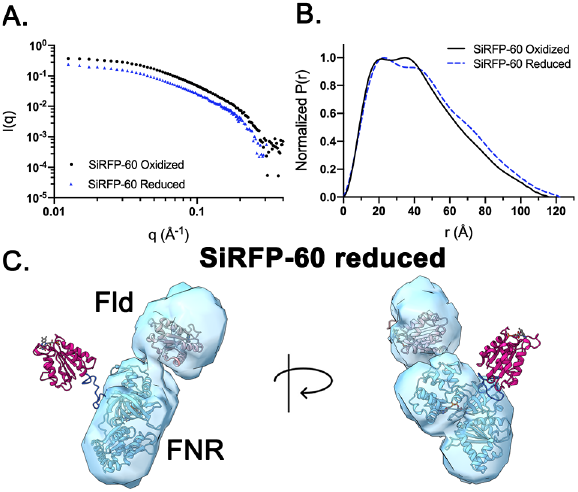
SiRFP-60 exhibits an extended conformation upon reduction. **A**. Scattering profile of oxidized (Tavolieri *et al*., 2019) and reduced SiRFP-60. **B**. The distance distribution plot of reduced SiRFP-60 shows an extended state and a shift in its bilobed peaks relative to the previously reported, oxidized form (Tavolieri *et al*., 2019). **C**. Model of reduced SiRFP-60 (blue) with the repositioned Fld domain from the crystal structure of SiRFP-60-Δ (light pink). (See **Figure 4D** for oxidized SiRFP-60).

## DISCUSSION

### SANS is the ideal technique to study the solution scattering of SiR

Here, we present a systematic analysis of dimeric SiR variants’ neutron scattering, allowing us to calculate the first solution structures of this essential metabolic oxidoreductase. By using SANS rather than SAXS, we were able to assess the contribution of each subunit to the scattering profiles because we could assemble dimers generated with hydrogenated SiRFP variants and deuterated SiRHP. Manipulation of the H_2_O/D_2_O ratios of the buffers then allowed us to isolate the scattering of SiRFP or SiRHP within the dimer. SANS is unique in allowing this type of contrast matching measurement, as is the fact that neutrons are less destructive than X-rays and therefore do not artificially impact the redox state of the system (Ankner *et al*, 2013). The series of structures show how SiR’s reductase and oxidase subunits interact and how the positioning of SiRFP’s Fld domain is impacted by both SiRHP binding and its redox environment.

The 8:4 stoichiometry of reductase to oxidase subunits in SiR has been long-hypothesized as important in its high-volume electron transfer chemistry by increasing the local concentration of reduced Fld domains for each SiRHP active site (Askenasy *et al*., 2018; Siegel & Davis, 1974; Tavolieri *et al*., 2019). This hypothesis predicts that SiRHP can recruit a transiently-interacting Fld domain from either a tightly-bound subunit *(cis* electron transfer) or an adjacent partner (*trans* electron transfer) (Figure 1D). SiR uses three NADPH molecules as reducing equivalents for each molecule of S^2-^ it generates, so access to more than one Fld domain would enhance the probability of a productive interaction with SiRHP. The changes that we see to the scattering envelopes from our SANS measurements on SiR dimer variants show for the first time how this might happen through motion of SiRFP’s Fld domain in response to both its oligomeric and redox states.

### SiRHP binding alters SiRFP-60’s interdomain orientation

When SiRFP is oxidized, it is in an open conformation where the small Fld domain is extended away from the curved FNR domain, connected by a 30 amino acid long linker (Figure 4D,Tavolieri *et al*., 2019). Upon SiRHP binding, the Fld domain rotates by about 30°relative to its position in the unbound form (Figure 3D), a conformational change that we confirmed with contrast matching experiments to isolate SiRFP-60’s contribution to the dimer’s scattering by matching-out D-SiRHP (Figure 4C). The resolution of the envelope is not sufficient to explain the mechanism by which SiRHP binding far from the Fld domain affects its conformation, but we are missing a very important detail about SiRHP: the structure of its N-terminus. As we have docked the atomic models, the N-terminal most amino acid in SiRHP’s structure (L81) is about 60 Å away from the last amino acid in SiRFP’s linker (P236). This distance could certainly be spanned by those N-terminal amino acids absent from current high-resolution structures (Crane *et al*., 1995) to affect the conformation of the linker and determine the position of the Fld domain relative to the FNR domain to which SiRHP binds.

### SiRHP undergoes compaction upon binding SiRFP-60

We also used SANS and contrast matching to reveal how SiRFP-60 binding affects the shape of SiRHP. SiRHP’s *D*_*max*_ decreases by almost 10 Å upon binding its partner, which can be visualized in the envelope function as a transition to a more globular shape. This compaction is likely the result of a rearrangement at its N-terminus, which is not resolved in the crystal structure, but known to be required for SiRFP binding (Askenasy *et al*., 2018; Crane *et al*., 1995). We propose that binding to SiRFP’s FNR domain either induces the flexible N-terminus to contract as it mediates intersubunit contacts or binds SiRFP as an extended peptide that would not contribute strongly to SiRHP’s scattering. That SiRHP binding affects SiRFP’s domain structure suggests that the latter may be occurring but the former cannot be excluded.

### SiRFP reduction repositions the Fld domain

In our model for the positions of SiRFP-60 and SiRHP within the envelopes modeled from neutron scattering, SiRHP is far from the Fld domain that would be responsible for funneling electrons to it within this minimal dimer (Figure 3D). This placement is consistent with the results from contrast matching experiments (Figures 4 and 5) as well as previously-reported mutational analysis that identified amino acids in the FNR domain of SiRFP as important for SiRHP binding (Askenasy *et al*., 2018; Askenasy *et al*., 2015). Nevertheless, *cis* electron transfer to a tightly-bound SiRHP in the minimal dimeric complex occurs, albeit at a rate of ∼50% that of the holoenzyme (Tavolieri *et al*., 2019; Zeghouf *et al*., 2000). Our SANS analysis reveals a dramatic reorientation of the Fld domain upon SiRFP-60 reduction (Figure 6).

## CONCLUSIONS

### Possible SiRFP conformations for *cis* or *trans* electron transfer

The envelope functions of SiR heterodimers determined from SANS provide the first solution structures depicting how SiRHP binds SiRFP’s FNR domain, far from the Fld domain from which the electrons move from the reductase to the oxidase. We confirmed our domain assignments by measuring scattering of SiR dimers containing a SiRFP variant that lacks the Fld domain as well as with the use of contrast variation to isolate each dimer component. Analyzing the hydrodynamic parameters and resulting envelope functions allowed us to make three observations that further our understanding of how electron transfer occurs in SiR. First, surprisingly, subunit binding impacts the position of the Fld domain even though it is far from where SiRHP binds, suggesting a mechanism by which *trans* electron transfer might take place in the context of the dodecameric holoenzyme because SiRHP binding positions the Fld domain away from it, perhaps pointing to an adjacent subunit (Figure 7A). Second, SANS of monomeric D-SiRHP and SiRFP/D-SiRHP heterodimers suggest SiRHP’s N-terminus undergoes a structural rearrangement upon complex assembly (Figure 7B). Third, superimposition of the bi-lobed feature of the reduced SiRFP-60 onto the envelope of the whole heterodimer shows that reduction swivels SiRFP-60’s Fld domain towards the binding position of SiRHP, suggesting a mechanism for *cis* electron transfer to a tightly-bound oxidase partner (Figure 7C). Together, these observations show that SiRFP’s interaction with SiRHP and redox state position SiRFP’s Fld domain for high-volume electron transfer between subunits.

**Figure 7:**
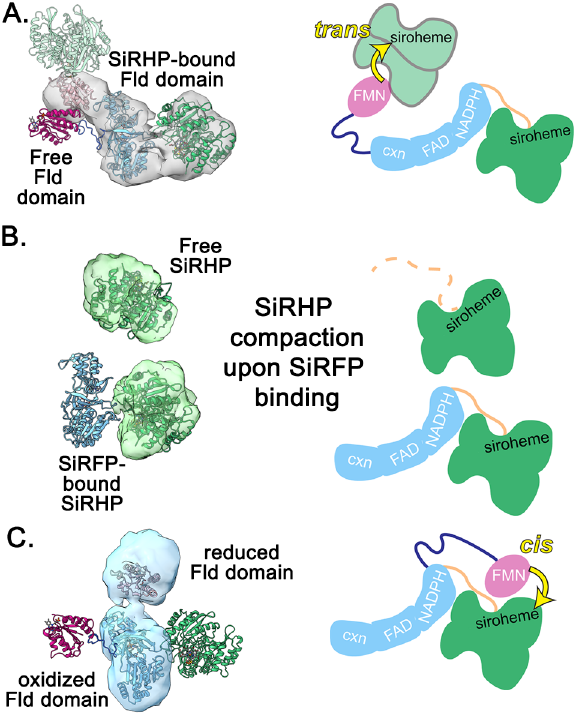
Model for SiRFP-SiRHP interactions. **A**. In the SiRFP-60/SiRHP dimer (gray density), SiRFP’s Fld domain (light pink) opens away from its FNR domain (light blue), perhaps positioning it for electron transfer to a SiRHP (transparent green) that is tightly-bound to an adjacent subunit. **B**. SiRHP (green density) becomes compact upon binding SiRFP’s FNR domain (light blue), likely from reorganization of its N-terminus. **C**. In reduced SiRFP-60, the Fld domain (light pink) is repositioned to orient towards SiRHP (green), suggesting providing a mechanism for electron transfer in *cis, i*.*e*. from SiRFP (light blue density) to a tightly-bound SiRHP (green).

## MATERIALS AND METHODS

### Expression, purification, and characterization of sulfite reductase proteins

Hydrogenated SiRFP and SiRHP proteins were expressed and purified as previously described (Askenasy *et al*., 2018; Askenasy *et al*., 2015; Tavolieri *et al*., 2019). Briefly, pBAD vectors (Thermo Fisher Scientific, Waltham, MA, USA) containing the genes encoding either N-terminally truncated/hexa-histidine tagged SiRFP (SiRFP-60) or untagged SiRHP from *Escherichia coli*(*E. coli*) were transformed into *E. coli* LMG194 cells (Invitrogen, Carlsbad, CA, USA) for recombinant protein expression. Proteins were purified to homogeneity with the use of nickel affinity, anion exchange, and size exclusion chromatography.

SiR heterodimers were formed by mixing purified monomeric SiRFP and SiRHP subunits followed by incubation for 30 minutes on ice. They were then loaded onto a 5 mL HisTrap FF nickel affinity chromatography column (Cytiva, Marlborough, MA, USA) that had been previously equilibrated with SPG buffer (17 mM succinic acid, 574 mM sodium dihydrogen phosphate, pH 6.8, 374 mM glycine, 200 mM NaCl) and eluted with a gradient of the same buffer containing 500 mM imidazole. Fractions obtained during the imidazole gradient were analyzed via SDS-PAGE and fractions containing heterodimer were loaded onto a Superose 6 10/300 size exclusion chromatography column (Cytiva, Marlborough, MA, USA) equilibrated with SANS buffer (50 mM KPi, pH 7.8, 100 mM NaCl, 1 mM EDTA) to separate residual monomers from the heterodimers.

Expression of D-SiRHP was carried out with fed-batch cultivation at Oak Ridge National Laboratory’s Bio-Deuteration Laboratory (ORNL, Oak Ridge, TN, USA). *E. coli* BL21 (DE3) cells (New England Biolabs, Ipswich, MA, USA) were transformed with the untagged SiRHP-expressing pBAD vector (Askenasy *et al*., 2015). Transformants were adapted to D_2_O by transferring an inoculum from Enfors minimal medium prepared with H_2_O and 100 μg/mL carbenicillin into the same medium with increasing D_2_O content (0, 50, and 70%) (Törnkvist, 1996). Once cells were growing in 70% D_2_O medium, a 250 mL preculture was used to inoculate 3.75 L of 70% D_2_O minimal media in a BioFlo 310 bioreactor (Eppendorf, Hauppauge, NY, USA). Cells were grown at 37 °C with dry, sterile air flow and agitation varying between 200 to 600 rpm to maintain dissolved oxygen above 30% saturation. 10% (w/v) NaOH in 70% D_2_O was fed on demand to maintain a pD > 7.3. Addition of feed solution consisting of 10% (w/v) H-glycerol, 0.2% MgSO_4_, 100 μg/mL carbenicillin in 70% D_2_O was initiated when the dissolved oxygen spiked upon depletion of H-glycerol from the batch medium. Fourteen hours post-inoculation and at an OD_600_ of 10, the temperature set point was reduced to 25 °C and D-SiRHP expression was induced by the addition of 0.05% L-arabinose in 70% D_2_O. 12 hours post-induction, cells were harvested by centrifugation at 6,000 x g for 40 minutes and resuspended in lysis buffer (65 mM KPi, pH 7.8, 200 mM NaCl, 1 mM EDTA) before being flash-frozen with liquid nitrogen (LN_2_).

D-SiRHP was purified as previously described for SiRHP with minor adjustments for scaling the amount of cell mass used during initial lysis and purification steps. Cell mass was thawed and diluted to 10% (w/v) with lysis buffer supplemented with 100 μg/mL PMSF and 1 μg/mL Pepstatin A. Cells were lysed at 15,000 psi in three passages through a water-cooled EmulsiFlex-C3 homogenizer (Avestin, Ottawa, ON, CA) and lysate collected in a beaker on ice. 0.1% (v/v) polyethyleneimine was added to the lysate while stirring for 30 minutes at 6 °C. Lysate was clarified by centrifugation at 13,000 x g for 25 minutes. Two successive ammonium sulfate cuts (31.5% and 52.5% (w/v)) were performed on the supernatant. The first precipitate was discarded after centrifugation at 13,000 x g for 25 minutes and the second was resuspended in lysis buffer. The resuspension was passed over a Sephadex G-25 desalting column and subsequently dialyzed into 5 mM KPi, pH 7.8, 1 mM EDTA solution overnight at 6 °C. The resulting solution was spun at 10,000 x g for 10 minutes before loading the supernatant onto a DEAE Sepharose Fast Flow anion exchange chromatography column (Cytiva, Marlborough, MA, USA) equilibrated with fresh dialysis buffer before being eluted with a gradient of 50 mM KPi, pH 7.8, 1 mM EDTA. Fractions containing D-SiRHP were concentrated with a 10 kDa MWCO polyethersulfone membrane-containing centrifugal concentrator and then loaded onto a Sephacryl S-300 HR size exclusion chromatography column (Cytiva, Marlborough, MA, USA) previously equilibrated with SANS buffer. The purified fractions were pooled and concentrated as before and then flash frozen with LN_2_. Partially deuterated SiR heterodimers were assembled in the same manner as their hydrogenated counterparts.

D-SiRHP was evaluated using small-angle X-ray scattering (SAXS) to assess deuterated sample quality in varying amounts of D_2_O on a Bio-SAXS System (Rigaku, Tokyo, Japan). SAXS was measured on D-SiRHP in SANS buffer composed of either 0, 75, or 100 % D_2_O. D-SiRHP’s scattering overlapped with that of hydrogenated SiRHP (H-SiRHP) under identical conditions and was free from aggregation (Figures S4A and B).

### Analytical ultracentrifugation

Sedimentation velocity experiments were performed using a ProteomeLab XL-1 analytical ultracentrifuge with the use of an AN60-Ti rotor and Epon-2 channel centerpieces (Beckman Coulter, Brea, CA, USA). Protein samples were diluted with SANS buffer to concentrations yielding an absorbance of 0.5 at 280 nm as measured with an 8454 UV–Vis spectrophotometer (Agilent, Santa Clara, CA, USA). Samples were then loaded into cell assemblies with sapphire windows for collecting absorbance data. Rotor speeds of 40,000 or 32,500 rpm were used for monomeric and heterodimeric SiR samples, respectively. All samples were run at 20 °C for 7 hours. Scan data were imported into UltraScan III software, after which 2-D spectrum and enhanced van Holde-Weischet analyses were completed to fit time-invariant noise and obtain sedimentation coefficient values, respectively (Demeler & Scott, 2005; Demeler & van Holde, 2004). Sedimentation coefficients were corrected for the density and viscosity of SANS buffer at 20 °C.

### Anaerobic reductions

SiRFP variants were reduced with sodium in an anaerobic glove box using SANS buffer prepared with 90% D_2_O that had been degassed using freeze-pump-thaw cycling and inert gas substitution. 10 molar equivalents of sodium dithionite were added to the sample and allowed to incubate for 30 minutes. In preparation for SANS measurements, 320 μL of reduced protein solution was loaded into 1 mm pathlength circular-shaped quartz cuvettes (Hellma USA, Plainville, NY, USA) and sealed with vacuum grease, rubber septa, and parafilm.

### SANS data collection

SANS data were collected on the Extended Q-range Small-Angle Neutron Scattering diffractometer (EQ-SANS, Beam Line 6) at the Spallation Neutron Source (SNS) located at ORNL. Two configurations were used in 60 Hz operation mode: 4 m sample-to-detector distance with 2.5-6.1 Å wavelength band and 1.3 m sample-to-detector distance with 4.0-8.3 Å wavelength band (Zhao *et al*, 2010) to obtain the relevant wavevector transfer, *Q*= 4π sin(θ)/λ, where 2θ is the scattering angle and λ is the neutron wavelength. Samples were loaded into 1 mm pathlength circular quartz cuvettes and data collected at 8 °C with the simultaneous introduction of dry air to prevent condensation on cuvettes. Scattering data were circularly averaged and reduced to one-dimensional scattering profiles using MantidPlot (Arnold *et al*, 2014). The measured scattering intensity was corrected for detector sensitivity and scattering contributions from buffers and empty cell, and then placed on absolute scale using a calibrated porasilica standard (Wignell & Bates, 1987). Replicate measurements were summed and those from each instrument configuration were merged. Incoherent background subtractions were also implemented in MantidPlot before the datasets were exported for analysis.

Hydrogenated samples were dialyzed into SANS buffer prepared with 90% D_2_O prior to SANS measurements to obtain sufficient contrast. Partially deuterated samples were dialyzed into mixtures containing 41 or 86% D_2_O to contrast match either the hydrogenated SiRFP or D-SiRHP of the heterodimer complexes, respectively. 41% D_2_O was chosen as the CMP for SiRFP from theoretical calculations based on amino acid sequence and agreement with CMPs of other hydrogenated proteins (A. E. Whitten 2008). The CMP of D-SiRHP (86% D_2_O) was determined experimentally by performing a contrast series of SANS measurements at 0, 41, 55, 75, 85, and 100% D_2_O, determining the forward scattering intensities *I(0)* of each measurement, and then plotting a linear fit of √*I(0)* vs. % D_2_O (Figures S4C and D).

For SANS measurements on reduced proteins, the anaerobicity of the samples were monitored over the course of data collection for changes in their scattering curves, indicative of changes due to re-oxidation. Comparisons of initial and final scattering curves showed no changes and the summed measurements were taken for analysis.

### SANS data analysis and modeling

After reduction in MantidPlot, data were imported into the ATSAS 3.0.1 suite (European Molecular Biology Laboratory, Hamburg Outstation), managed in PRIMUS (Konarev *et al*, 2003; Petoukhov *et al*, 2012). *R*_*g*_ and *I(0)* were determined in PRIMUS, whereas *D*_*max*_ and *P(r)* analyses were calculated in GNOM (Svergun, 1992). *R*_*g*_ and *I(0)* were obtained using a Guinier approximation (Guinier, 1955) (a linear fit of a ln[*I(q)*] vs. *q*^*2*^ plot, where *R*_*g*_ x *q*_*max*_ < 1.3). *D*_*max*_ values were obtained through an indirect Fourier transform of the scattering data, yielding real-space distance distribution functions for each specimen, with *D*_*max*_ determined by where the function decays to *r*= 0. The *P(r)* data were used as inputs for *ab initio* modeling using DENSS (DENsity from Solution Scattering) software to create density maps that were then refined to produce the final scattering envelopes (Grant, 2018). Comparisons of SANS data to theoretical scattering curves were calculated in CRYSON (Svergun *et al*, 1998) with protein chain deuteration fraction and D_2_O fraction in the solvent specified to mimic the experimental contrast conditions. High-resolution structures obtained from the Protein Data Bank (PDB) were positioned into solution structures in UCSF ChimeraX (Berman *et al*, 2000; Goddard *et al*., 2018).

## AUTHOR CONTRIBUTIONS

D.T.M., C.B.S., G.N., and M.E.S. performed SANS experiments and analyzed data. D.T.M. performed wet lab experiments and analyzed data. D.T.M. and M.E.S. designed wet lab experiments. D.T.M and K.L.W. designed and performed deuteration experiments. M.E.S. designed the project and wrote the manuscript together with D.T.M. All authors reviewed, approved, and contributed to the final version of the manuscript.

## ACKNOWLEDGEMENTS

We thank Claudius Mundoma for helpful conversations regarding the analysis of AUC data, Christopher Stroupe for comments on the manuscript, and Tristan Dilbeck for careful reading of the manuscript. A portion of this research at ORNL’s Spallation Neutron Source was sponsored by the Scientific User Facilities Division, Office of Basic Energy Sciences, U.S. Department of Energy. The Office of Biological and Environmental Research also supported work at the Oak Ridge National Laboratory Center for Structural Molecular Biology. This work was supported by National Science Foundation grants MCB1856502 and CHE1904612 to M.E.S.

## FIGURE LEGENDS

**Figure S1:**
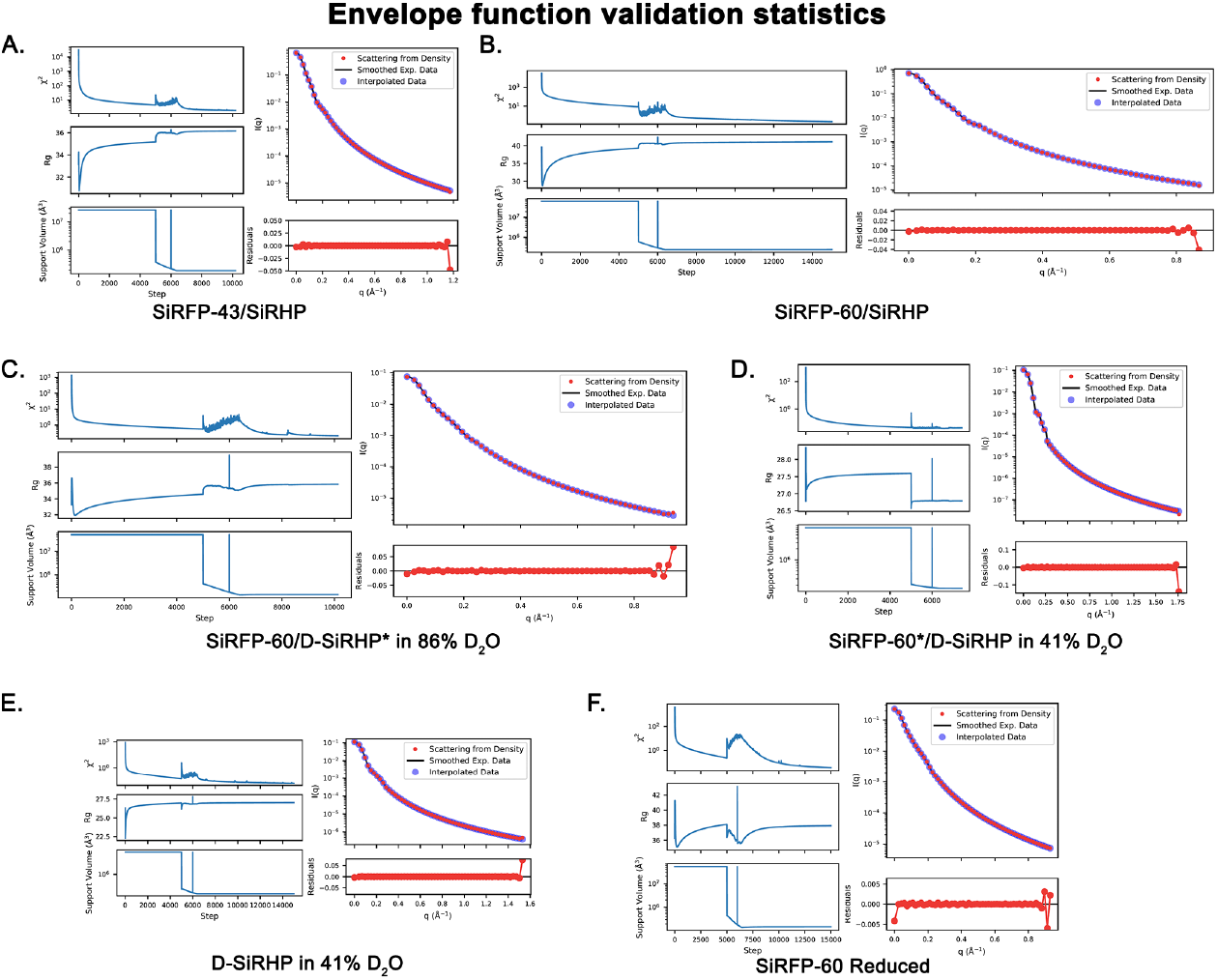
Theoretical scattering calculated from *ab initio* models fit their experimental data. For each panel, plots on the left side of each panel show *X*^*2*^ (top), *R*_*g*_(middle), and support volume (bottom) converging through the course of modeling steps. The top right plot shows scattering from the modeled density fit to its experimental data and the bottom right shows their residuals. Aberrant residual points at high *q*, and sometimes low *q*, are outside of the *q* range used in the modeling. Datasets used as inputs for modeling were truncated to remove aberrant data points at low *q* due to proximity with the neutron beam and at high *q* to remove noisy incoherent scattering. **A**. SiRFP-43/SiRHP **B**. SiRFP-60/SiRHP **C**. SiRFP-60/D-SiRHP* **D**. SiRFP-60*/SiRHP **E**. D-SiRHP **F**. SiRFP-60 (reduced).

**Figure S2:**
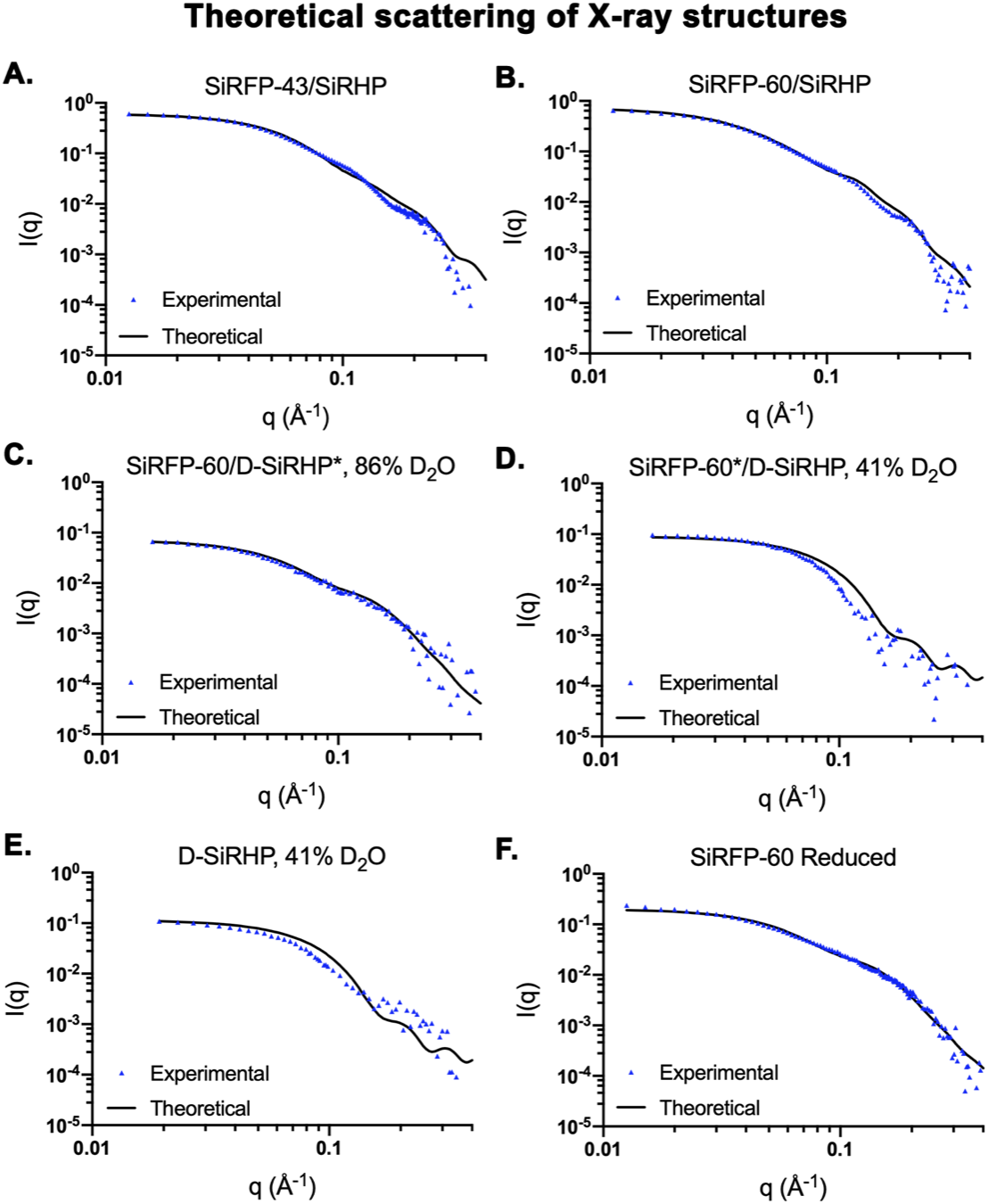
Theoretical scattering curves calculated from the positioned X-ray crystal structures reflect the experimental data, except in cases where experimental solution scattering conditions imparted changes relative to static crystal structures. **A**. SiRFP-43/SiRHP **B**. SiRFP-60/SiRHP **C**. SiRFP-60/D-SiRHP* **D**. SiRFP-60*/SiRHP **E**. D-SiRHP **F**. SiRFP-60 (reduced).

**Figure S3:**
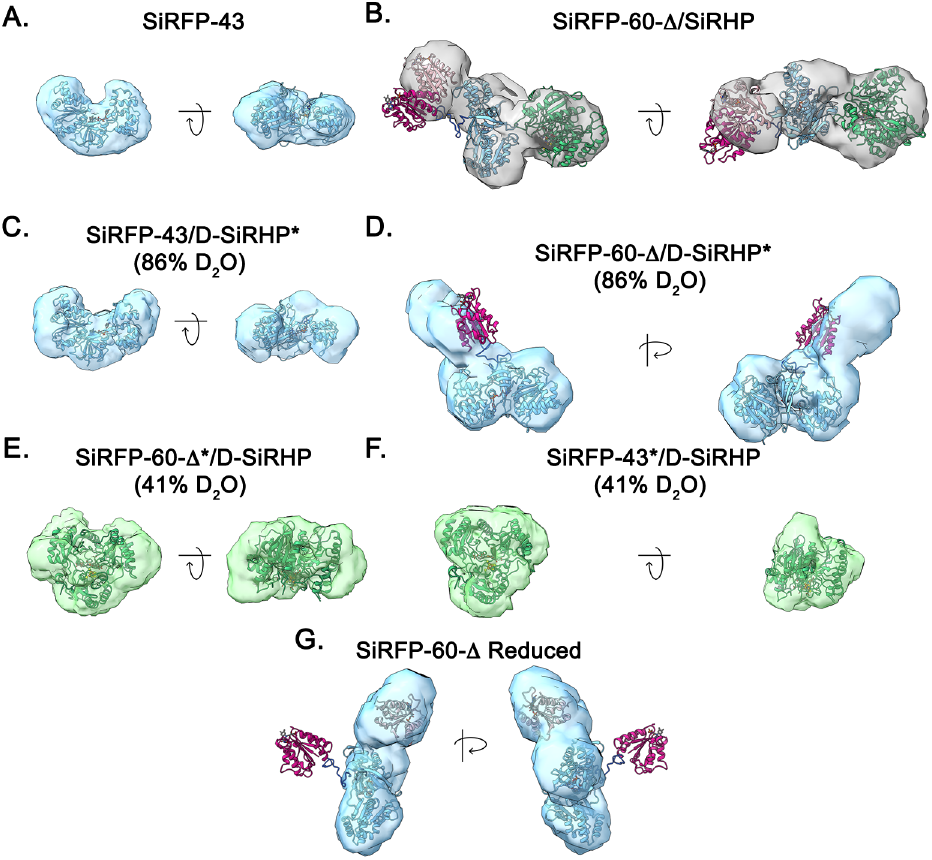
*Ab initio* models of control specimen. Domains and proteins are colored as in the main text (SiRFP’s FNR domain = light blue, SiRFP-60-Δ’s Fld domain = dark pink, placed Fld domain = light pink, SiRHP = green) **A**. SiRFP-43 **B**. SiRFP-60-Δ/SiRHP **C**. SiRFP-43/D-SiRHP* **D**. SiRFP-60-Δ/SiRHP* **E**. SiRFP-60-Δ*/D-SiRHP **F**. SiRFP-43**\*/**D-SiRHP **G**. SiRFP-60-Δ (reduced).

**Figure S4:**
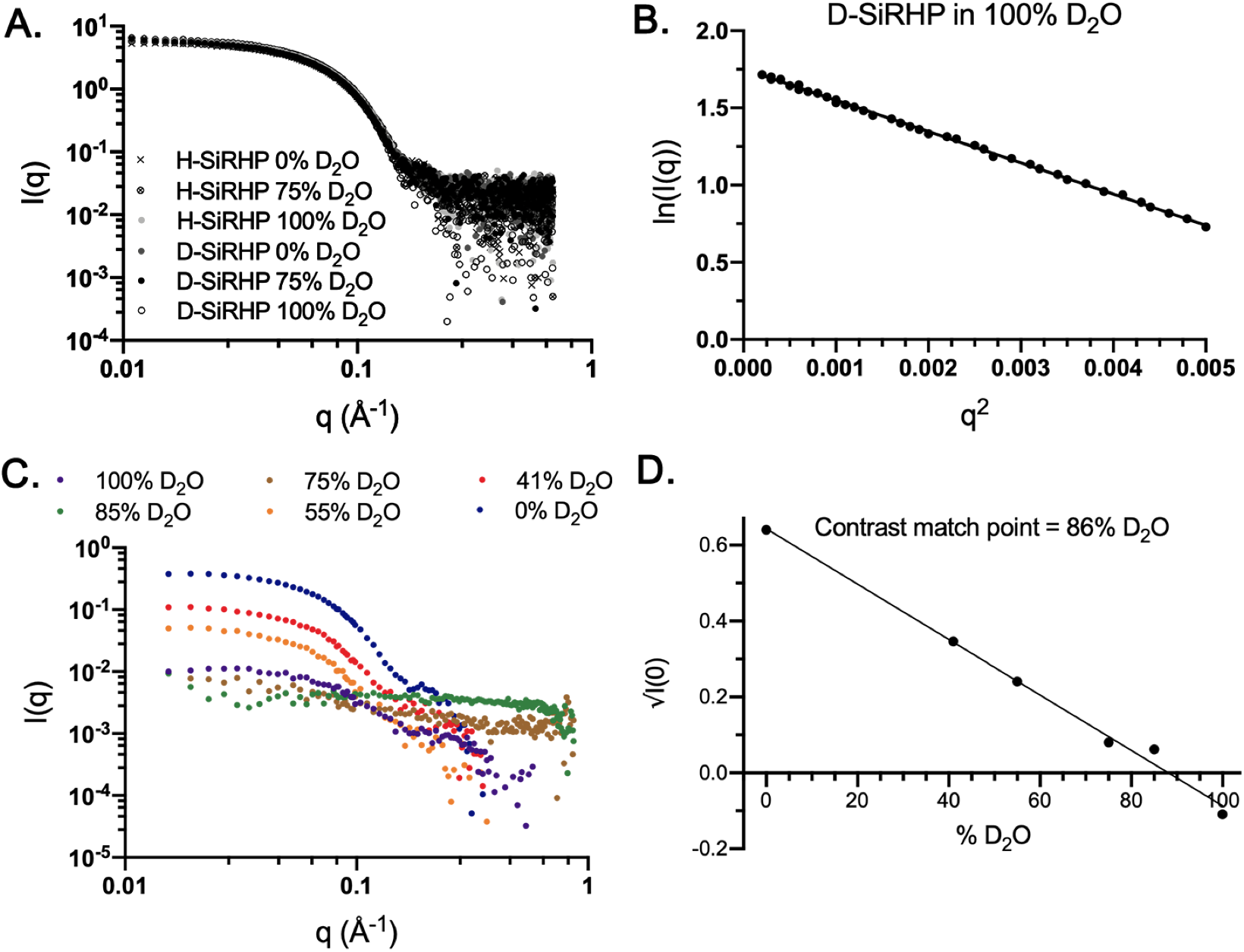
Characterization of D-SiRHP. **A**. SAXS profiles of D-SiRHP in various H_2_O/D_2_O mixtures overlap with those of H-SiRHP. **B**. Guinier analysis from SAXS of D-SiRHP in 100% D_2_O shows a monodispersed sample. **C**. Contrast variation SANS of D-SiRHP in six different percentages of D_2_O shows a decrease in signal intensity, *I(q)*, as the contrast match point is approached (see panel for key). **D**. The X-intercept of a linear fit to √*I(0)* vs % D_2_O indicates D-SiRHP’s match point, 86% D_2_O.

